# Deep Learning Predicts Survival Across Squamous Tumor Entities From H&E Stains: Insights from Head and Neck, Esophagus, Lung and Cervical Cancer

**DOI:** 10.1101/2025.03.31.646351

**Authors:** Verena Bitto, Xiaofeng Jiang, Michael Baumann, Jakob Nikolas Kather, Ina Kurth

## Abstract

Computational pathology-based models are becoming increasingly popular for extracting biomarkers from images of cancer tissue. However, their validity is often only demonstrated on a single unseen validation cohort, limiting insights into their generalizability and posing challenges for explainability. In this study, we developed models to predict overall survival using haematoxylin and eosin (H&E) slides from FFPE samples in head and neck squamous cell carcinoma (HNSCC). By validating our models across diverse squamous tumor entities, including, head and neck (HR = 1.58, 95% CI = 1.17–2.12, p = 0.003), esophageal (non-significant), lung (HR = 1.31, 95% CI = 1.13–1.52, p < 0.001) and cervical (HR = 1.39, 95% CI = 1.10–1.75, p = 0.005) squamous cell carcinomas, we showed that the predicted risk score captures relevant information for survival beyond HNSCC. Correlation analysis indicated that the predicted risk score is strongly associated with various clinical factors, including human papillomavirus status, tumor volume and smoking history, although the specific factors vary across cohorts. These results emphasize the necessity for comprehensive validation and in-depth assessment of computational pathology-based models to better characterize the underlying patterns they learn during training.

## Introduction

Survival models estimate the likelihood of a specific clinical endpoint occurring in a patient in the future by incorporating clinical characteristics (such as age and sex) or biomedical markers (from laboratory assays or images)^1^. The field of Computational Pathology (CP) has proven capable of extracting such prognostic markers directly from routinely available haematoxylin and eosin (H&E)-stained whole-slide images (WSIs)^2^. More recently, it was shown that a single label per whole-slide image (WSI), can be sufficient to train prognostic models for diverse cancer entities through so-called multiple instance learning (MIL)^3^. This is typically achieved by first extracting lower dimensional embeddings from patches of WSIs using existing pre-trained models^3^. Such models, when trained on a broad and diverse range of data, are typically referred to as foundation models^4^. To date, it has been shown that CP models can generalize effectively to multicentre setups^5^. We argue here that, if these models learn characteristics inherently related to the tumor morphology, they should also possess the capability to generalize across tumors of the same histological type originating from different locations. In this study, we developed and trained a model to predict survival directly from pathological slides of patients with head and neck squamous cell carcinoma (HNSCC) and evaluated its prognostic value further in patients suffering from squamous tumor entities of the esophagus, lung and cervix.

HNSCC refers to a group of cancers originating from distinct anatomical sites, namely the oral cavity, the nasopharynx, the oropharynx, the hypopharynx, and the larynx^6^, which can be further split by their histology^7^. All of these cancers originate from squamous epithelial tissue. Other squamous-originating tumor entities include esophageal cancer (ESCC), cervical squamous cell carcinoma (CSCC) and lung squamous cell carcinoma (LSCC)^8^. One common risk factor for certain squamous cell carcinomas is the presence of persistent virus infection with the human papillomavirus (HPV), predominantly oropharynx cancer and CSCCs^9^. The evidence of an association between HPV and other squamous cancers, such as ESCC^10^ and LSCC^11^, is less consistent. Naturally, certain risk factors for squamous carcinomas apply to specific anatomical sites. For example, alcohol consumption is associated with carcinomas in the oral cavity, pharynx, larynx and esophagus, tissues that are directly exposed^8^. These shared characteristics, along with their distinct differences, make SCCs particularly compelling for study within a CP framework.

We investigated whether the MIL of features extracted from H&E stains using foundation models can directly predict survival in patients with squamous cell carcinomas, when trained on only one particular tumor entity. First, we trained a model on H&E stained sections from the publicly available TCGA HNSC cohort. Second, we evaluated whether the results are transferable to a multi-center cohort of patients with locally advanced HNSCC, treated with a standardized, state-of-the-art radiochemotherapy (RCTx) protocol. Patients treated with primary RCTx usually present with a more advanced-stage disease compared to those treated with surgery, leading to poorer overall survival. Thus, it is of significant importance to assess whether the trained models can identify high-risk candidates within a well-defined treatment modality. Lastly, we investigated whether the results generalize to squamous cell carcinomas of other tumor entities, i.e., CESC, ESCC and LSCC, from TCGA.

## Materials and Methods

### Patient Cohorts

Histology slides and clinical data of three HNSCC cohorts were utilized to train, validate and test the model. For training, all 472 diagnostic slides from 450 cases of the TCGA-HNSC^12^ cohort were downloaded from the GDC portal, covering diverse head and neck anatomical sites and patient treatment regimes. Cohorts of a single centre (73 slides of 73 patients, Dresden, hereafter referred to as BU, with patients receiving primary radio[chemo]therapy) and a multi-centre study (157 slides of 156 patients, DKTK-ROG receiving primary RCTx^13^) were used as validation and test cohorts. For testing the transferability to other squamous cell carcinoma entities, H&E stains of CESC (project id TCGA-CESC, 269 cases, 279 diagnostic slides), ESCC (project id TCGA-ESCA, 156 cases, 158 diagnostic slides), and LSCC (project id TCGA-LUSC, 478 cases, 512 diagnostic slides), of TCGA were downloaded from the GDC portal.

For TCGA cohorts, clinical data were taken from the TCGA Pan-Cancer Clinical Data Resource (TCGA-CDR)^14^. The HPV infection status, assessed using RNA-Seq and exome sequencing, and smoking status were obtained for all TCGA cohorts from the study conducted by Campbell et al^15^. If a major HPV type was assigned in Campbell et al., the different types were further summarized into two categories according to Rade et.^16^, as follows: Category 1 summarizes phylogenic clades with A9 being comprised of HPV 16, 31, 33, 35, 52, 58, 67, and A7 being comprised of HPV 18, 39, 45, 59, 68, 70, 85, and 97. Category 2 clusters HPV 16-related (HPV 16, 31, and 52) or HPV 18-related (HPV 18 and 45) subtypes. For the BU and DKTK HNSCC cohorts, HPV status was determined by p16 immunohistochemical staining (p16) and HPV type 16 (HPV16) DNA. To account for the different assessment methods, all figures and tables clearly distinguish between ‘HPV status’ for TCGA, and ‘p16’ or ‘HPV16’ for BU and DKTK. Categories with many different levels, such as Pathological T stage, were consolidated into broader groups (e.g., Stage I for Stage IA and Stage 1B).

### Image Processing and Deep Learning Techniques

#### Data Preprocessing

Slides from DKTK were primarily scanned with a ZEISS Axioscan 7 at 20x magnification, and the resulting CZI files were converted to a TIFF format supported by OpenSlide. H&E stains from TCGA were utilized without further initial steps. All slides were pre-processed using the corresponding step of the STAMP pipeline^17^. Briefly, STAMP splits the H&E slide into patches, discards background and noisy patches and extracts histological feature vectors by utilizing foundation models. We utilized the built-in support of the foundation models CTransPath^18^, UNI^19^ and CONCH^20^.

### Model Development

For training, the extracted histological features of patches from the 450 TCGA HNSC patients were fed into a MIL model to optimize survival risk scores using a Cox partial likelihood loss described previously by Jiang et al.^5^. Unlike their model, our model was trained solely on the endpoint of overall survival (OS) rather than a combination of two endpoints. This approach was chosen because, apart from OS, the endpoints of TCGA and BU/DKTK were closely linked but not identical, e.g., disease-free interval (TCGA) versus loco-regional control (BU/DKTK). Additionally, only 134 patients out of the 450 TCGA HNSC patients had available follow-up information for disease-free interval.

An Adam optimizer with an initial learning rate of 1e-3, a batch size of 16, and weights for L1 and L2 regularization of 0.01 were used for training. For UNI and CONCH, the configuration was adapted to a learning rate of 1e-5 and a batch size of 32. The models were trained for 50 epochs, with early stopping set to 10 epochs to prevent overfitting. The best models were selected based on its ability to correctly rank survival outcomes in the validation cohort, as measured using the concordance index.

For validation, all slides and patients of BU HNSCC and 43 slides of 42 patients of two centres from DKTK-ROG HNSCC were used, referred to thereafter as BU/DKTKmber. This resulted in a total of 115 individuals in the validation cohort. For testing on the same tumor entity, slides of the 114 patients from the remaining centres of DKTK-ROG HNSCC were processed, referred to thereafter as DKTKrem. For testing on other squamous tumor entities, the individual performance of the model on TCGA cohorts of CESC, ESCC and LSCC was evaluated.

### Visualization

Attention heatmaps and corresponding representative tiles were generated as described in Jiang et al.^5^ for the validation cohorts where the predicted risk scores demonstrated prognostic value for survial.

### Experimental Design

The predicted risk scores for overall survival were evaluated as input for a Cox regression model to determine if this covariate is indicative of actual overall survival. Multivariable Cox regression was performed on the HNSCC validation cohort (DKTKrem), as risk factors had previously been identified specifically for this cohort^13^. Apart from Cox analysis, patients were grouped into risk groups (high, low, medium) according to their risk core. For the HNSCC cohorts (TCGA HNSC, BU/DKTKmber, DKTKrem) the risk groups were defined based on the 25th and 75th percentiles of the risk scores from the training set. For the other cohorts (TCGA ESCA, TCGA LUSC, TCGA CESC), risk groups were assigned based on the 25th and 75th percentiles of the respective cohort. This approach was chosen due to the noticeable shift in distribution when the model is applied to cohorts with different tumor entities. High- and low-risk groups were evaluated using Kaplan-Meier estimates, and statistical significance was assessed through log-rank tests. Differences in characteristics of high- and low risk groups, were assessed as follows: For all categorical variables a Pearson’s Chi-squared test was performed and a ‘Missing’ level was added for testing if more than five individuals had missing values for a category. For all continuous variables a Wilcoxon rank-sum test was conducted. For all statistical tests, a p-value < 0.05 was considered statistically significant.

Feature extraction of histological slides and model training/evaluation was performed using Python (3.11.11) by means of STAMP^17^ and marugoto^5^ with torch (2.6.0). All downstream analyses were conducted using R (4.4.0) and the following packages: The overall performance per cohort was evaluated using the Harrell’s concordance index (C-Index) via the survcomp^21^ (1.54.0) package. For visualization of survival curves, computation of log-rank tests and Cox regression models, including hazard ratios (HR) and their confidence intervals (CI), the survminer^22^ (0.5.0), ggplot2^23^ (3.5.1), and survival^24^ (3.7–0) packages were utilized. Characteristics of risk groups were compared using tableone^25^.

### Data and Code availability

Extraction of histological features from H&E stains was performed using STAMP^17^ (https://github.com/KatherLab/STAMP/). The model was trained by means of survival branch of marugoto (https://github.com/KatherLab/marugoto/tree/survival) as initially described in Jiang et al.^5^

## Results

We trained histopathology-based models to predict survival in patients with squamous cell carcinomas (HNSCC, ESCA, LUSC, CESC) by leveraging different foundation models to extract features with the training solely been based on the TCGA HNSC cohort (**Figure 1**). The predicted risk score, an aggregate of the risk scores of all H&E patches of a patient, is used as continuous variable in univariable Cox regression analysis and to group individuals into risk groups (low, medium, high) for performing Kaplan-Meier estimates and log-rank tests.

**Figure 1.**
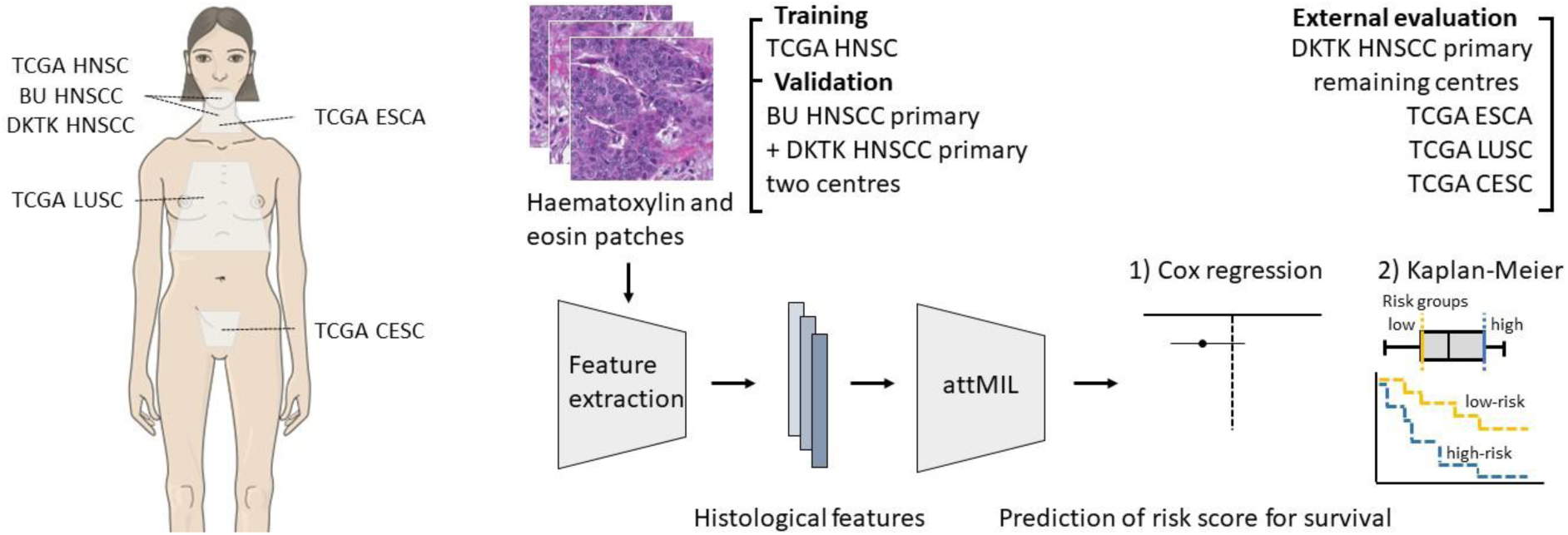
Analysis workflow to predict overall survival in squamous cell carcinomas. Left: Squamous cell carcinomas under investigation. Cohorts include TCGA head and neck squamous cell carcinoma (TCGA HNSC), DKTK head and neck squamous cell carcinoma (DKTK HNSCC), BU head and neck squamous cell carcinoma (BU HNSCC), TCGA esophageal carcinoma (TCGA ESCA), TCGA lung squamous cell carcinoma (TCGA LUSC) and TCGA cervical squamous cell carcinoma and endocervical adenocarcinoma (TCGA CESC). Right: Patches of haematoxylin and eosin (H&E) stains are processed by extracting histological features by means of foundation models. These features are then used to train a model to predict survival based on the TCGA HNSC cohorts. The external evaluation includes cohorts from the same tumor entity and cohorts of other squamous cell carcinomas.

### Prognostic value in HNSCC cohorts

**Table 1** shows the performance of the training, validation and test cohorts when the feature extraction was carried out with CTransPath, indicating that the predicted risk score can prognosticate survival in HNSCC. A similar pattern emerges if only high- and low-risk patients are evaluated using Kaplan Meier estimates (**Figure 2**), although the corresponding log-rank test for the validation cohort showed no significant difference. Interestingly, when examining whether high- and low-risk patients share common characteristics, different aspects emerged for each cohort (**Supplementary Table 1**) According to the training cohort, low-risk patients were more likely to be HPV positive and/or reformed smokers, both factors known to have a positive effect on survival^26, 27^. Counterintuitively, they also had a higher clinical stage compared to the high-risk group, but no significant difference in the pathological T stage, pointing to the multifactorial nature of cancer prognosis. Finally, low-risk patients had a significant smaller number of slide patches available, referred to as the technical variable NPATCH. The difference in NPATCH suggests that patients with more tissue collected were also at higher risk of death. These correlations, however, were only partly reflected in the HNSCC test cohort. Here, patients at higher risk also exhibited a higher tumor volume. Notably, in the training cohort, only the corresponding pathological T stage was a known parameter. In line with the potential association with HPV status observed in the training cohort, all 12 patients in the high-risk group in the test cohort were HPV-negative (based on p16 and HPV16 testing). Drawing on insights from previous studies on the DKTK HNSCC cohort, multivariable Cox regression was performed. **Table 2** highlights that among N stage, p16, the logarithm of the primary tumor volume [ln(GTV)] and the predicted risk score, only the tumor volume remains statistically significant in its association with overall survival. However, the hazard ratios suggest that the risk score may offer some independent prognostic value. To investigate this further, we created a combined factor of the risk score and the GTV, which categorizes their levels into three groups: “High” when both GTV and risk group are high, “medium” for any combination of high, medium, or low levels between the two factors, and “low” when both GTV and risk group are low. **Figure 3** shows that the combination of our risk score and GTV stratifies patients by survival, despite the remaining small sample size.

**Figure 2.**
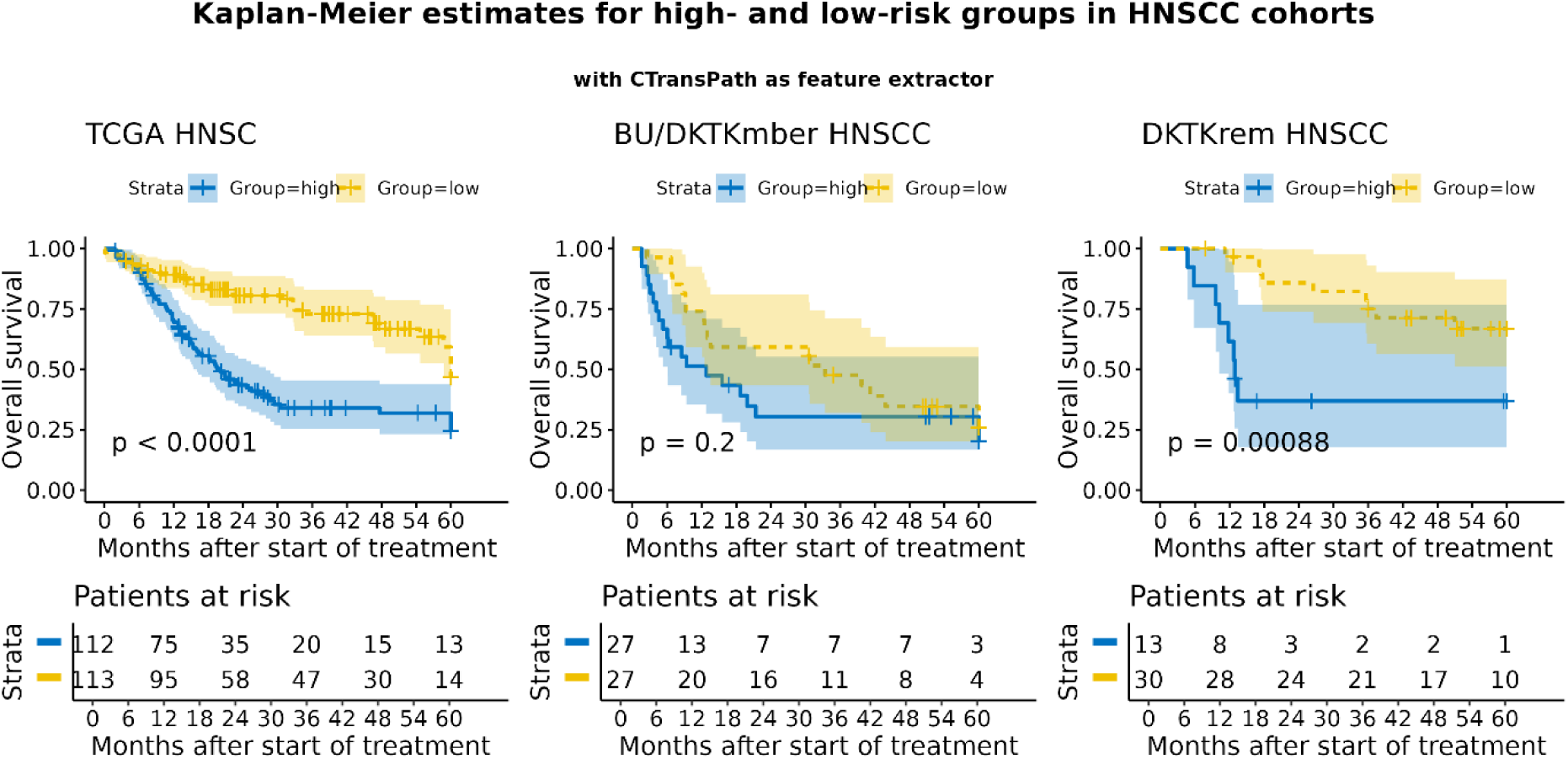
Survival analysis on head and neck squamous cell carcinoma (HNSCC) cohorts using CTransPath as feature extractor. Kaplan-Meier estimates of high and low risk group for overall survival in training cohort (left, TCGA HNSC), validation cohorts (middle, BU HNSCC and DKTK HNSCC) and test cohort (right, DKTK Subset HNSCC).

**Figure 3.**
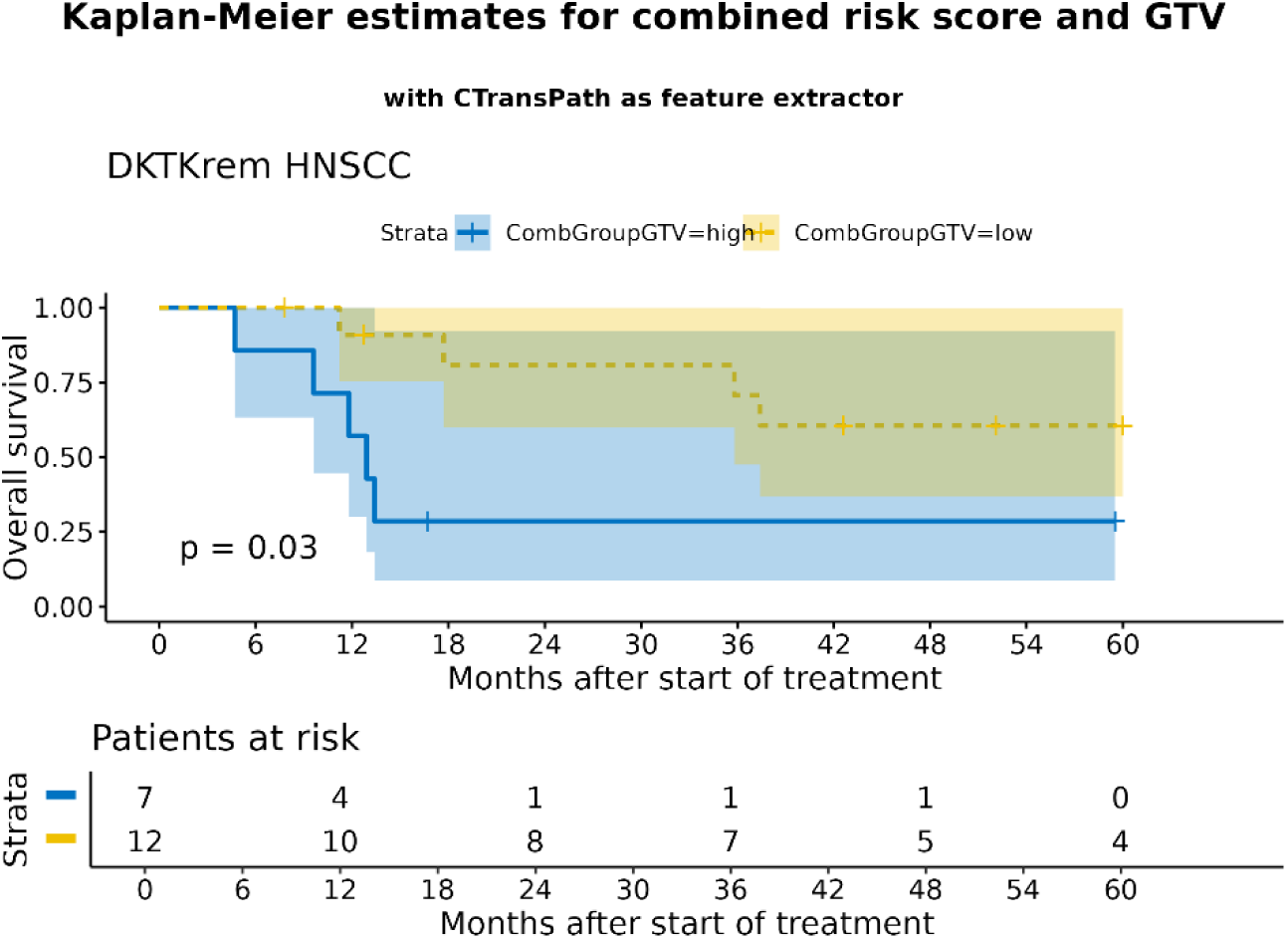
Survival analysis on HNSCC test cohort using a combined factor of the risk score and GTV. The combined factor categorizes the levels of GTV and risk group into three groups: “high” when both GTV and risk group are high, “medium” for any combination of high, medium, or low levels between the two factors, and “low” when both GTV and risk group are low.

**Table 1.**
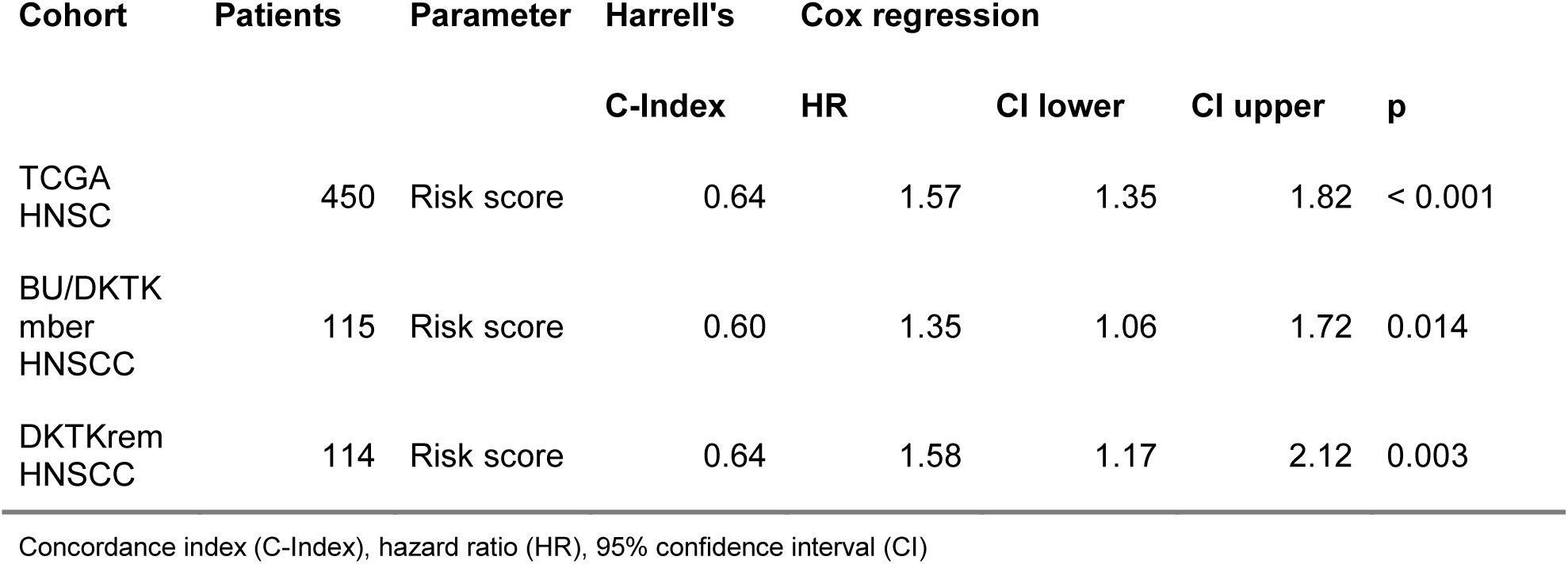
Prognostic value of predicted risk score for overall survival in head and neck squamous cell carcinoma (HNSCC) cohorts when features were extracted with CTransPath. Shown are the Harrell’s concordance index and hazard ratios from univariable Cox regression analysis based on predicted risk scores considering all patients of the individual cohorts.

**Table 2.**
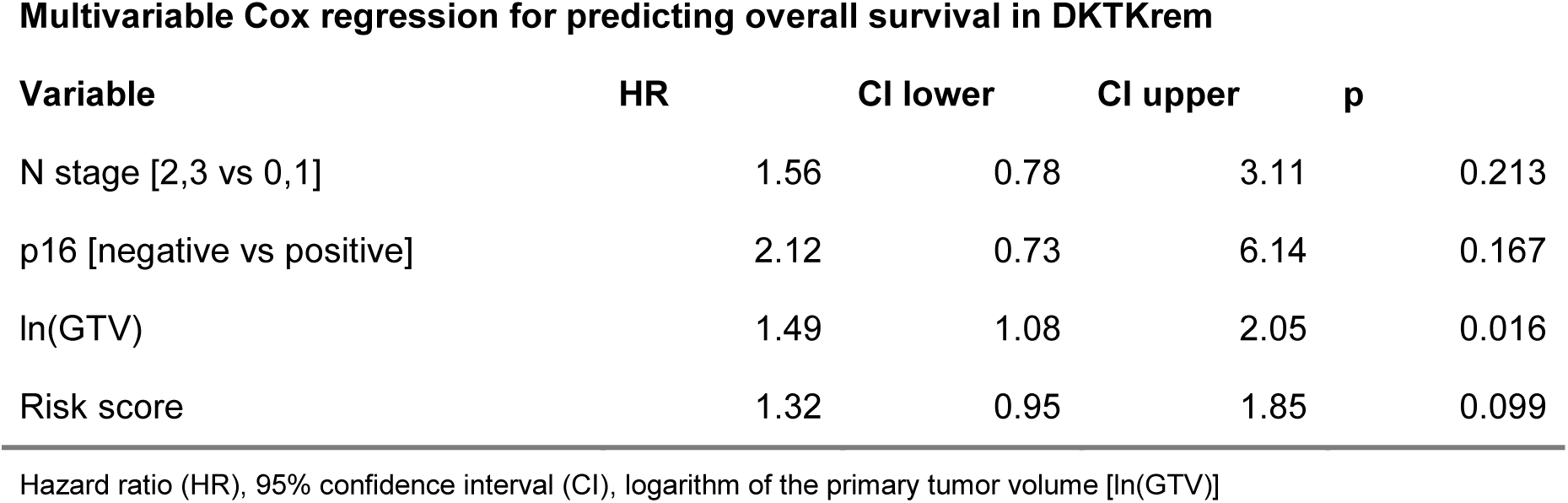
Multivariable Cox regression analysis of DKTKrem HNSCC cohort with features extracted using CTransPath.

**Multivariable Cox regression for predicting overall survival in DKTKrem**
| Variable | HR | CI lower | CI upper | p |
| --- | --- | --- | --- | --- |
| N stage [2,3 vs 0,1] | 1.56 | 0.78 | 3.11 | 0.213 |
| p16 [negative vs positive] | 2.12 | 0.73 | 6.14 | 0.167 |
| ln(GTV) | 1.49 | 1.08 | 2.05 | 0.016 |
| Risk score | 1.32 | 0.95 | 1.85 | 0.099 |

Hazard ratio (HR), 95% confidence interval (CI), logarithm of the primary tumor volume [ln(GTV)]

### Prognostic value in other squamous cohorts

To evaluate whether our trained model extends its prognostic value across squamous carcinomas, we examined its performance on TCGA ESCA, TCGA LUSC and TCGA CESC. While the predicted risk score was not prognostic for esophageal carcinomas, which are anatomically adjacent to the neck, it showed an association for survival in patients with LUSC and CESC (**Table 3**). The clinical relevance of the risk score was further supported when splitting patients into high- and low-risk group patients to fit Kaplan-Meier estimates (**Figure 4**). Exploration of common characteristics in TCGA LUSC and CESC revealed that a greater number of elderly individuals (age > 65) were categorized into the high-risk group in the TCA LUSC cohort. For TCGA CESC, lifetime non-smokers were at lower risk for death, which shows consistency with literature indicating that smoking is associated with unfavorable disease free and overall survival outcomes in cervical cancer patients^28^. Interestingly, the model appears to separate patients by histology type in CESC (i.e., squamous, adenosquamous and adenocarcinoma), with more individuals with squamous histology classified as low-risk. No relevant difference in HPV status was observed in any of the three cohorts, except TCGA CESC, where the significant difference cannot be reliably interpreted due to the large number of unclassified patients.

**Figure 4.**
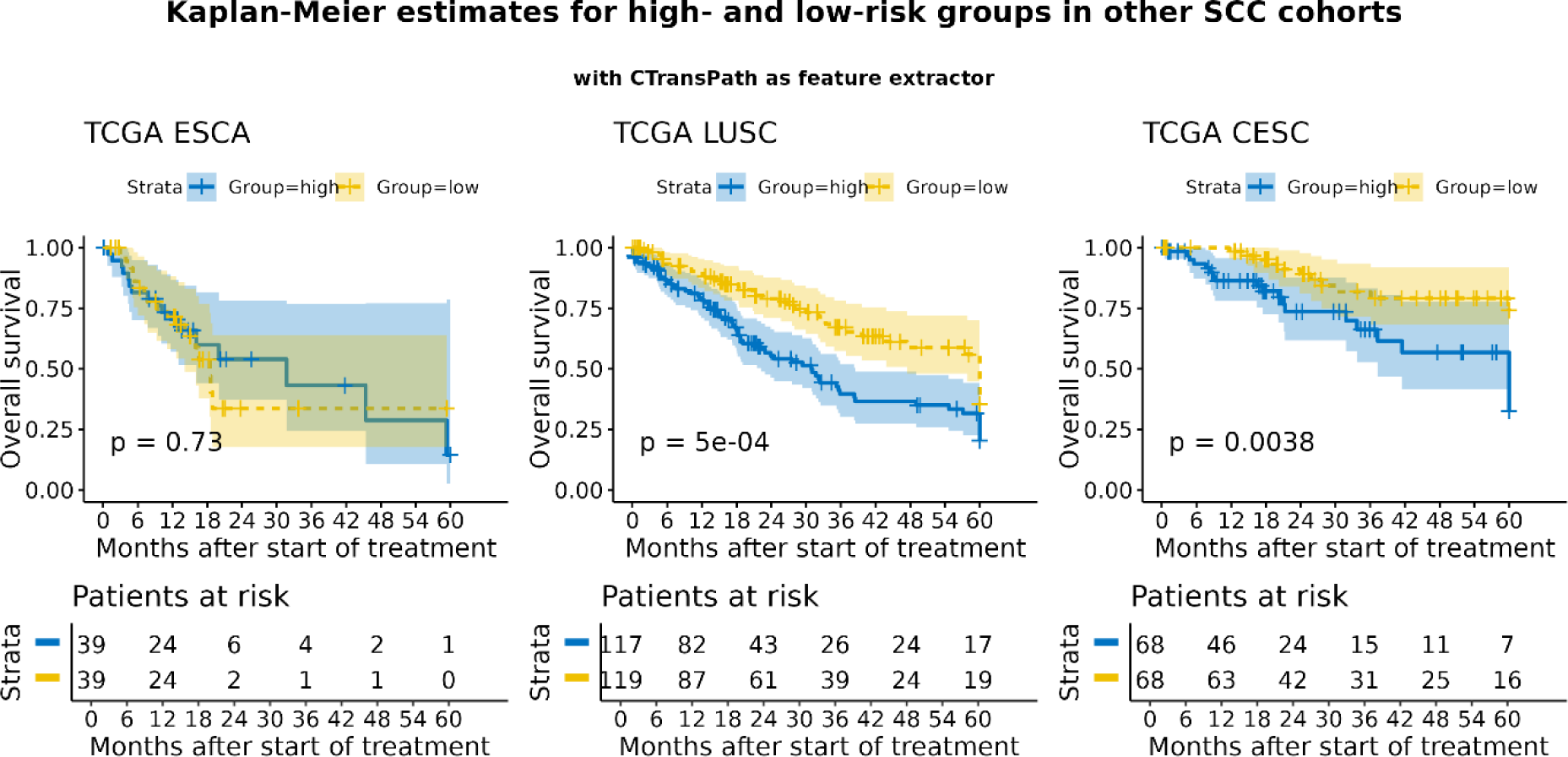
Survival analysis on squamous cell carcinoma cohorts using CTransPath as feature extractor. Kaplan-Meier estimates of high and low risk group for overall survival in TCGA esophageal carcinoma (TCGA ESCA, left), TCGA lung squamous cell carcinoma (TCGA LUSC, middle) and TCGA cervical squamous cell carcinoma and endocervical adenocarcinoma (TCGA CESC, right).

**Table 3.** Prognostic value of predicted risk score for overall survival in other squamous cell carcinoma cohorts when features were extracted with CTransPath. Shown are the Harrell’s concordance index and hazard ratios from univariable Cox regression analysis based on predicted risk scores considering all patients of the individual cohorts.

| Cohort | Patients | Parameter | Harrell's | Cox regression |  |  |  |
| --- | --- | --- | --- | --- | --- | --- | --- |
|  |  |  | C-Index | HR | CI lower | CI upper | p |
| TCGA ESCA | 156 | Risk score | 0.50 | 0.96 | 0.74 | 1.24 | 0.741 |
| TCGA LUSC | 477 | Risk score | 0.59 | 1.31 | 1.13 | 1.52 | < 0.001 |
| TCGA CESC | 269 | Risk score | 0.60 | 1.39 | 1.10 | 1.75 | 0.005 |

Concordance index (C-Index), hazard ratio (HR), 95% confidence interval (CI)

### Visualization of attention maps

To identify which areas within H&E stains were most relevant for ranking in the survival models, attention heatmaps and representative tiles are visualized in **Supplementary Figure 1**. In all cases shown, the representative tile contain visible tumor cells. Notably, tiles containing no tissue or those with noise, e.g., markers, were removed before training using the STAMP framework, as indicated by the black background in these regions.

### Evaluation of different foundation models

To assess if the results shown are specific to the use of CTransPath as feature extractor, we investigated the performance of models built on the foundation models CONCH and UNI. **Table 4** and **Table 5** show similar prognostic value when using CONCH as a feature extractor for the overall performance across all patients and cohorts, compared to CTransPath. This is further supported by the corresponding Kaplan-Meier estimates of high- and low risk groups (**Supplementary Figure 2** and **Supplementary Figure 3**). Like with the CTransPath-based model, patients categorized into the low-risk groups had significantly better survival outcome in TCGA HNSC, DKTKrem HNSCC, TCGA LUSC and TCGA CESC when features were extracted with CONCH. Since the UNI-based model showed no prognostic value in the HNSCC test cohort (results not shown), we did not evaluate it on other squamous entities.

**Table 4.** Prognostic value of predicted risk score for overall survival in head and neck squamous cell carcinoma (HNSCC) cohorts when features were extracted with CONCH. Shown are the Harrell’s concordance index and hazard ratios from univariable Cox regression analysis based on predicted risk scores considering all patients of the individual cohorts.

| Cohort | Patients | Parameter | Harrell's | Cox regression |  |  |  |
| --- | --- | --- | --- | --- | --- | --- | --- |
|  |  |  | C-Index | HR | CI lower | CI upper | p |
| TCGA<br>HNSC | 450 | Risk score | 0.59 | 1.33 | 1.15 | 1.54 | < 0.001 |
| BU/DKTK<br>mber<br>HNSCC | 115 | Risk score | 0.59 | 1.34 | 1.06 | 1.71 | 0.016 |
| DKTKrem<br>HNSCC | 114 | Risk score | 0.62 | 1.39 | 1.05 | 1.82 | 0.019 |

Concordance index (C-Index), hazard ratio (HR), 95% confidence interval (CI)

**Table 5.** Prognostic value of predicted risk score for overall survival in other squamous cell carcinoma cohorts when features were extracted with CONCH. Shown are the Harrell’s concordance index and hazard ratios from univariable Cox regression analysis based on predicted risk scores considering all patients of the individual cohorts.

| Cohort | Patients | Parameter | Harrell's | Cox regression |  |  |  |
| --- | --- | --- | --- | --- | --- | --- | --- |
|  |  |  | C-Index | HR | CI lower | CI upper | p |
| TCGA<br>ESCA | 156 | Risk score | 0.57 | 1.09 | 0.86 | 1.39 | 0.456 |
| TCGA<br>LUSC | 477 | Risk score | 0.56 | 1.17 | 1.03 | 1.34 | 0.019 |
| TCGA<br>CESC | 269 | Risk score | 0.62 | 1.32 | 1.04 | 1.67 | 0.021 |

Concordance index (C-Index), hazard ratio (HR), 95% confidence interval (CI)

## Discussion

Our study investigated if CP models are able to predict survival directly from pathological slides in squamous tumor entities using MIL and foundation models as feature extractors. We showed that, although trained on a small cohort of TCGA HNSC (450 individuals) and selected based on our HNSCC BU/DKTKmber validation cohort, the models generalized well to unseen patients with locally advanced HNSCC. Additionally, we showcased that their prognostic value is not limited to HNSCC but extends to other squamous tumor entities, demonstrated in cohorts from TCGA LUSC and TCGA CESC. Although SCCs classified by their site of origin, different studies have found that they share common molecular properties^15, 29^, some of which manifest in histopathological slides. We argue that validating findings from CP models on a range of similar tumor entities is crucial for a more comprehensive understanding of the learned characteristics. If we would have limited our analysis to HNSCC, one would likely conclude that our model picks up primarily signals of HPV infection, suggested by the results of the training and validation cohort (see **Supplementary Table** 1). However, the model also stratified patients according to high and low risk in TCGA LUSC, in which 96% of all patients are HPV negative (**Table 6**). Consequently, the risk score must pick up also signals from other characteristics visible in histopathological slides. This aligns with the understanding that cancer is a multifactorial disease influenced by various genetic and environmental factors. For example, Mulder et al. found that HNSCC in non-smokers and non-drinkers is more often linked to HPV and p16 overexpression, while in a third of HPV-negative tumors, TP53 mutations show a mutational profile associated with aging and ultraviolet light exposure rather than tobacco consumption^30^. Apart from predicting clinical outcomes, one key question is whether models trained on histological features offer prognostic value beyond established clinical factors. To address that question, we performed multivariable Cox regression analysis for survival, incorporating risk factors that have been identified in previous publications for the DKTK cohort13. **Table 2** suggests that the predicted risk score carries prognostic value (p value 0.099) besides the GTV. This becomes more evident when combining both the risk score and GTV into one factor, allowing for clear stratification of patients by survival (**Figure 3**). Considering that the risk score correlates with several other covariates (i.e., p16 and ln[GTV], **Supplementary Table 1**), this could obscure its true prognostic value both in the Cox model and in the combined factor. Further exploration of the connection to GTV could enhance the explainability of CP-based models; however, this factor is often missing in clinical cohorts, as GTV estimates typically require additional imaging modalities like CT scans. Given that GTV is a known independent risk factor for survival – along with loco-regional control and other endpoints – in various squamous cell carcinomas^31–34^, it is striking that it is not commonly reported in studies.

**Table 6.** Patients characteristics of TCGA esophageal carcinoma (TCGA ESCA), TCGA lung squamous cell carcinoma (TCGA LUSC) and TCGA cervical squamous cell carcinoma and endocervical adenocarcinoma (TCGA CESC).

|  |  | **TCGA CESC**<br>**(N=269)** | **TCGA ESCA**<br>**(N=156)** | **TCGA LUSC**<br>**(N=477)** | **Overall**<br>**(N=902)** |
| --- | --- | --- | --- | --- | --- |
| Age | <=65 | 240 (89.2%) | 106 (67.9%) | 179 (37.5%) | 525 (58.2%) |
|  | >65 | 29 (10.8%) | 50 (32.1%) | 289 (60.6%) | 368 (40.8%) |
|  | Missing |  |  | 9 (1.9%) | 9 (1.0%) |
| Gender | Female | 269 (100.0%) | 21 (13.5%) | 121 (25.4%) | 411 (45.6%) |
|  | Male |  | 135 (86.5%) | 356 (74.6%) | 491 (54.4%) |
| HPV_status | Negative | 9 (3.3%) | 89 (57.1%) | 458 (96.0%) | 556 (61.6%) |
|  | Positive | 211 (78.4%) |  | 3 (0.6%) | 214 (23.7%) |
|  | Missing | 49 (18.2%) | 67 (42.9%) | 16 (3.4%) | 132 (14.6%) |
| Smoking | Current | 52 (19.3%) | 29 (18.6%) | 121 (25.4%) | 202 (22.4%) |
|  | Never | 108 (40.1%) | 31 (19.9%) | 17 (3.6%) | 156 (17.3%) |
|  | Reformed (Any) | 45 (16.7%) | 27 (17.3%) | 317 (66.5%) | 389 (43.1%) |
|  | Missing | 64 (23.8%) | 69 (44.2%) | 22 (4.6%) | 155 (17.2%) |
| Clinical_stage | Stage I | 36 (13.4%) | 3 (1.9%) |  | 39 (4.3%) |
|  | Stage II | 50 (18.6%) | 29 (18.6%) |  | 79 (8.8%) |
|  | Stage III | 42 (15.6%) | 20 (12.8%) |  | 62 (6.9%) |
|  | Stage IV | 20 (7.4%) | 10 (6.4%) |  | 30 (3.3%) |
|  | Missing | 121 (45.0%) | 94 (60.3%) | 477 (100.0%) | 692 (76.7%) |
| Pathological_T_<br>stage | Stage I/II |  | 75 (48.1%) | 383 (80.3%) | 458 (50.8%) |
|  | Stage III/IV |  | 45 (28.8%) | 87 (18.2%) | 132 (14.6%) |
|  | Missing | 269 (100.0%) | 36 (23.1%) | 7 (1.5%) | 312 (34.6%) |

|  |  | TCGA CESC<br>(N=269) | TCGA ESCA<br>(N=156) | TCGA LUSC<br>(N=477) | Overall<br>(N=902) |
| --- | --- | --- | --- | --- | --- |
| HPV_clade | A9 | 155 (57.6%) |  | 2 (0.4%) | 157 (17.4%) |
|  | A7 | 50 (18.6%) |  |  | 50 (5.5%) |
|  | other | 6 (2.2%) |  |  | 6 (0.7%) |
|  | Missing | 58 (21.6%) | 156 (100.0%) | 475 (99.6%) | 689 (76.4%) |
| HPV_related | HPV 16-related | 137 (50.9%) |  | 2 (0.4%) | 139 (15.4%) |
|  | HPV 18-related | 43 (16.0%) |  |  | 43 (4.8%) |
|  | Missing | 89 (33.1%) | 156 (100.0%) | 475 (99.6%) | 720 (79.8%) |

Although our models showed prognostic value for survival in squamous cell carcinomas, a fixed parameter setup can limit model generalization. As illustrated by the technical variable NPATCH in **Supplementary Table 1**, patients with more tissue samples available were more often grouped into the high-risk group during training. Considering that the risk score is an aggregate of the risk scores from a bag, i.e., a concept in MIL describing a fixed-size subset of patches, the bag size may introduce potential model bias. Such technical bias would imply that the model consistently assigns lower scores to patients with statistically fewer H&E patches than the pre-defined bag size. Alternatively, the findings might represent a real correlation between the number of tissue collected from a patient and their risk of death. For instance, NPATCH may correlate with a true prognostic factor, such as the corresponding tumor volume in a patient. This is consistent with the results of the DKTKrem cohort, in which high-risk patients had a larger tumor volume and a higher mean NPATCH value (**Supplementary Table 1**) compared to their low-risk counterparts. Overall, there appears to be little evidence for a technical bias, which is further supported by the results from TCGA LUSC where high-risk coincides with a considerably lower number of patches available compared to the low-risk group (**Supplementary Table 2**).

Apart from the need of hyperparameter optimization in CP models, the sample size plays an essential role. Although training was performed on a small sample size, our model generalized well to other cohorts and is expected to improve as more training/validation data is added. To the best of our knowledge, we are the first, which evaluated the prognostic value of CP models on diverse squamous tumor entities. Our workflow is based on the approach outlined in Jiang et al.^5^, in which a model was trained to predict survival in colorectal cancer by considering overall survival and disease-free survival. For HNSCC, CP models have so far been centered on predicting specific clinical characteristics rather than clinical endpoints. Among the first, previous work of Kather et al. indicated that HPV and Epstein-Barr-Virus infection can be predicted through histopathology-based deep learning in patients with HNSCC and gastric cancer, respectively^35^. Later, Klein et al. trained a model on patients with oropharyngeal squamous carcinomas to predict their HPV status^36^. Rather than using the complete H&E stains of each patient as input, they performed semantic segmentation of viable tumor regions before training. Similarly, Wang et al. proposed a HPV score for patients with HNSCC by utilizing MIL without requiring tumor region annotations^37^. Both HPV prediction models were trained based on ResNet architectures rather than relying on the feature extraction capabilities of foundation models. Later, Wang et al. also established a molecular subtyping score for cervical squamous cell carcinoma based on MIL of different cohorts using UNI as feature extractor^38^.

Foundation models have become more and more popular in CP, making the selection of the most appropriate model increasingly challenging. Recently published examples include UNI2^39^, Virchow2^40^, and Prov-GigaPath^41^. Considering the small amount of data available for training, we opted for foundation models with a relatively low number of embeddings, i.e., 768 in CTransPath, 512 in CONCH and 1024 in UNI. Newer models typically come with a larger number of embeddings, such as 1280 embeddings in case of Virchow2 and 1536 in ProvGigaPath. Recently, Neidlinger et al. showed that different foundation models focus on different areas in the tissue^42^. Adding to this, Alfasly et al. highlighted that foundation models like UNI lack retrieval performance when it comes to extracting features from heterogeneous tissue like in cervical cancer^43^. This might explain why we receive better results for predicting survival when using CTransPath and CONCH over UNI as foundation model. Notably, CTransPath is pretrained on TCGA data, which may give it an advantage over other foundation models in extracting relevant histological features. However, we also showed that we receive similar results when using CONCH for feature extraction, which was trained on proprietary datasets rather than on slides from TCGA^20^. Additionally, our results on the unseen HNSCC cohort DKTKrem indicate that the findings are not limited to specific TCGA cohorts but also generalize to other proprietary datasets. Rather than emphasizing the superiority of one foundation model over another, our study highlights the need to expand explainability methods across tumor entities in CP-based models.

## Conclusion

In conclusion, we showcased that our CP-based models predict survival directly from pathological slides in HNSCC and other squamous tumor entities. In view of all the presented results, CP survival models likely pick up signals related to a diverse set of clinical-related factors such as HPV status, histological subtypes and smoking status. The findings highlight the need for broader evaluation of CP models on diverse tumor entities in order to improve explainability.

## Ethics statement

Ethical approval for the multicentre retrospective analyses of clinical and imaging data was obtained from the Ethics Committee at the Technische Universität Dresden, Germany (EK397102014) and from the Ethics Committees of all DKTK partner sites. The requirement for individual informed consent was waived owing to the retrospective nature of the study. All methods were performed in accordance with the relevant guidelines and regulations. Follow-up data of patients were collected using the RadPlanBio Platform^44^ at the DKTK partner site Dresden.

## Acknowledgements

We thank Dr. Annett Linge and Prof. Dr. Steffen Löck for their guidance and insightful discussions on findings in the BU/DKTK cohorts and Rosemarie Euler-Lange for her excellent technical assistance and support in digitizing the H&E stains.

## Author contributions

Conceptualization: VB, JK

Data Curation: VB

Formal Analysis: VB

Investigation: VB

Methodology: XJ

Project administration: JK, IK

Supervision: JK, IK, MB

Validation: VB

Visualization: XJ

Writing – original draft: VB

Writing – review & editing: VB, XJ, JK, IK, MB

All authors read and approved the final manuscript.

---

**Supplementary Figure 1.**
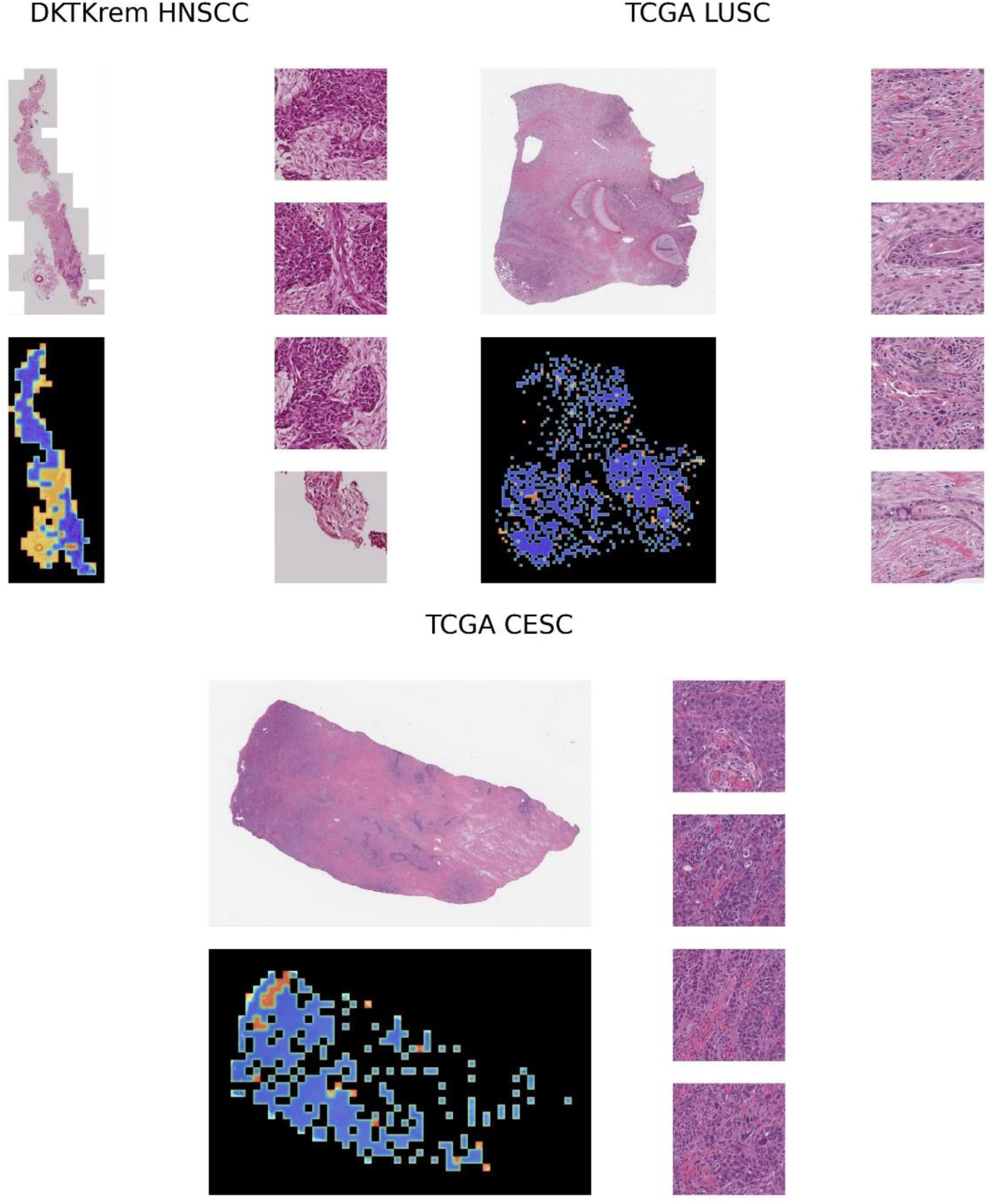
Exemplary images of haematoxylin and eosin stains and attention heatmaps from patients of DKTKrem HNSCC (upper, left), TCGA LUSC (upper, right) and TCA CESC (bottom). Displayed are the original H&E stain (top left), the attention heatmap (bottom, left) as outputted by the CTransPath-based model, where important regions appear in shades of red and less relevant areas in shades of blue, and representative tiles (right) extracted from the red-highlighted regions.

**Supplementary Figure 2.**
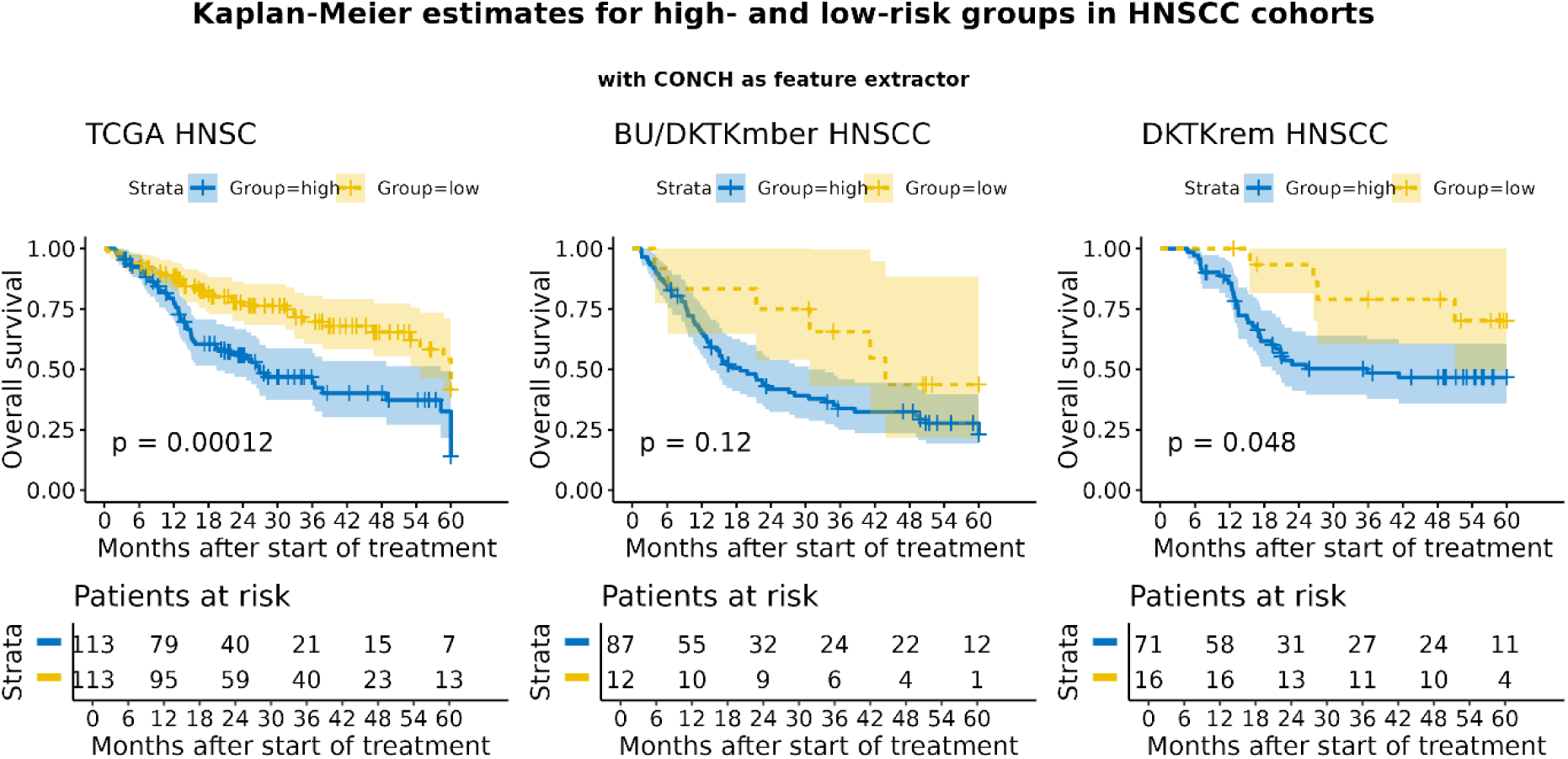
Survival analysis on head and neck squamous cell carcinoma (HNSCC) cohorts using CONCH as feature extractor. Kaplan-Meier estimates of high and low risk group for overall survival in training cohort (left, TCGA HNSC), validation cohorts (middle, BU HNSCC and DKTK HNSCC) and test cohort (right, DKTK Subset HNSCC).

**Supplementary Figure 3.**
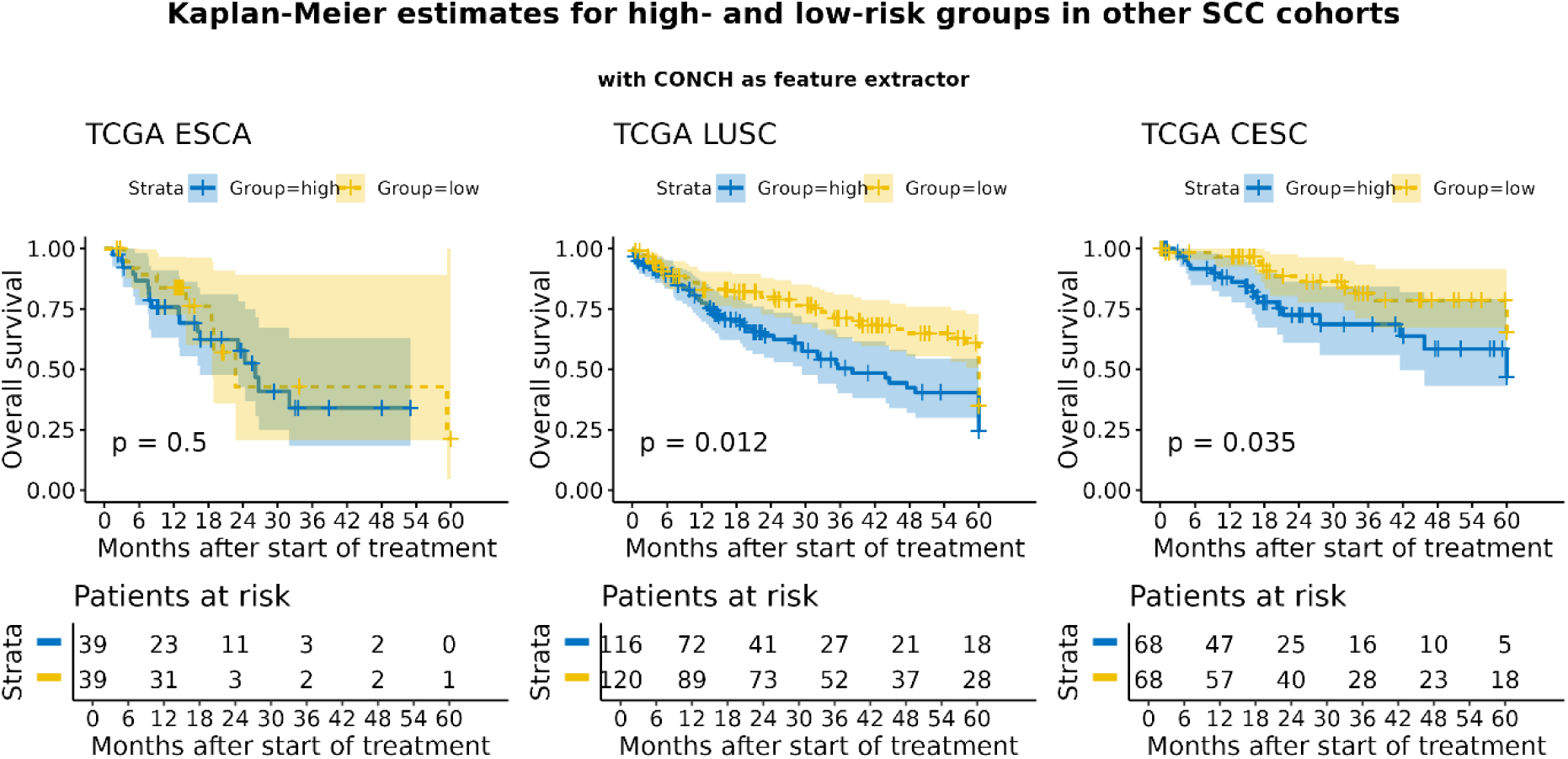
Survival analysis on squamous cell carcinoma cohorts using CONCH as feature extractor. Kaplan-Meier estimates of high and low risk group for overall survival in TCGA esophageal carcinoma (TCGA ESCA, left), TCGA lung squamous cell carcinoma (TCGA LUSC, middle) and TCGA cervical squamous cell carcinoma and endocervical adenocarcinoma (TCGA CESC, right).

Supplementary Table 1 Shared characteristics among high- and low risk groups in head and neck squamous cell carcinoma (HNSCC) cohorts. Empty entries indicate that the specific variable was not available for the cohort under investigation. [See excel file]

Supplementary Table 2 Shared characteristics among high- and low risk groups in other squamous cell carcinoma (SCC) cohorts. [See excel file]

## References

1. Uncategorized References

1. Collins GS, Reitsma JB, Altman DG, Moons KG. Transparent reporting of a multivariable prediction model for individual prognosis or diagnosis (TRIPOD): the TRIPOD Statement. BMC Med. 2015;13:1. 10.1186/s12916-014-0241-z.

2. Echle A, Rindtorff NT, Brinker TJ, Luedde T, Pearson AT, Kather JN. Deep learning in cancer pathology: a new generation of clinical biomarkers. Br J Cancer. 2020;124(4):686–696. 10.1038/s41416-020-01122-x.

3. Song AH, Jaume G, Williamson DFK, et al. Artificial intelligence for digital and computational pathology. Nature Reviews Bioengineering. 2023;1(12):930–949.

4. Bommasani R, Hudson DA, Adeli E, et al. On the Opportunities and Risks of Foundation Models. 2021.

5. Jiang X, Hoffmeister M, Brenner H, et al. End-to-end prognostication in colorectal cancer by deep learning: a retrospective, multicentre study. Lancet Digit Health. 2024;6(1):e33–e43. 10.1016/S2589-7500(23)00208-X.

6. Johnson DE, Burtness B, Leemans CR, Lui VWY, Bauman JE, Grandis JR. Head and neck squamous cell carcinoma. Nature Reviews Disease Primers. 2020;6(1).

7. Woolgar JA, Triantafyllou A. Squamous cell carcinoma and precursor lesions: clinical pathology. Periodontol 2000. 2011;57(1):51–72. 10.1111/j.1600-0757.2011.00389.x.

8. Yan W WI, Emmert-Buck MR, Erickson HS. Squamous Cell Carcinoma - Similarities and Differences among Anatomical Sites. Am J Cancer Res. 2011;1(3):275–300.

9. Egawa N. Papillomaviruses and cancer: commonalities and differences in HPV carcinogenesis at different sites of the body. Int J Clin Oncol. 2023;28(8):956–964. 10.1007/s10147-023-02340-y.

10. Petrelli F, De Santi G, Rampulla V, et al. Human papillomavirus (HPV) types 16 and 18 infection and esophageal squamous cell carcinoma: a systematic review and meta-analysis. J Cancer Res Clin Oncol. 2021;147(10):3011–3023. 10.1007/s00432-021-03738-9.

11. Karnosky J, Dietmaier W, Knuettel H, et al. HPV and lung cancer: A systematic review and meta-analysis. Cancer Rep (Hoboken). 2021;4(4):e1350. 10.1002/cnr2.1350.

12. The Cancer Genome Atlas Network. Comprehensive genomic characterization of head and neck squamous cell carcinomas. Nature. 2015;517(7536):576–582.

13. Linge A, Lohaus F, Löck S, et al. HPV status, cancer stem cell marker expression, hypoxia gene signatures and tumour volume identify good prognosis subgroups in patients with HNSCC after primary radiochemotherapy: A multicentre retrospective study of the German Cancer Consortium Radiation Oncology Group (DKTK-ROG). Radiother Oncol. 2016;121(3):364–373.

14. Liu J, Lichtenberg T, Hoadley KA, et al. An Integrated TCGA Pan-Cancer Clinical Data Resource to Drive High-Quality Survival Outcome Analytics. Cell. 2018;173(2):400–416 e411. 10.1016/j.cell.2018.02.052.

15. Campbell JD, Yau C, Bowlby R, et al. Genomic, Pathway Network, and Immunologic Features Distinguishing Squamous Carcinomas. Cell Rep. 2018;23(1):194–212 e196. 10.1016/j.celrep.2018.03.063.

16. Rader JS, Tsaih SW, Fullin D, et al. Genetic variations in human papillomavirus and cervical cancer outcomes. Int J Cancer. 2019;144(9):2206–2214. 10.1002/ijc.32038.

17. El Nahhas OSM, van Treeck M, Wolflein G, et al. From whole-slide image to biomarker prediction: end-to-end weakly supervised deep learning in computational pathology. Nat Protoc. 2025;20(1):293–316. 10.1038/s41596-024-01047-2.

18. Wang X, Yang S, Zhang J, et al. Transformer-based unsupervised contrastive learning for histopathological image classification. Med Image Anal. 2022;81:102559. 10.1016/j.media.2022.102559.

19. Chen RJ, Ding T, Lu MY, et al. Towards a general-purpose foundation model for computational pathology. Nature Medicine. 2024;30(3):850–862.

20. Lu MY, Chen B, Williamson DFK, et al. A visual-language foundation model for computational pathology. Nat Med. 2024;30(3):863–874. 10.1038/s41591-024-02856-4.

21. Schroder MS, Culhane AC, Quackenbush J, Haibe-Kains B. survcomp: an R/Bioconductor package for performance assessment and comparison of survival models. Bioinformatics. 2011;27(22):3206–3208. 10.1093/bioinformatics/btr511.

22. Kassambara A, Kosinski M, Biecek P. survminer Drawing Survival Curves using ‘ggplot2’; 2024. https://github.com/kassambara/survminer.

23. Wickham H. ggplot2: Elegant Graphics for Data Analysis; 2016. https://ggplot2.tidyverse.org.

24. Therneau TM. A Package for Survival Analysis in R; 2024. https://CRAN.R-project.org/package=survival.

25. Yoshida K. tableone: R package to create “Table 1”, description of baseline characteristics with or without propensity score weighting; 2022. Accessed 2025/01/31 0.13.2. https://github.com/kaz-yos/tableone.

26. O’Rorke MA, Ellison MV, Murray LJ, Moran M, James J, Anderson LA. Human papillomavirus related head and neck cancer survival: a systematic review and meta-analysis. Oral Oncol. 2012;48(12):1191–1201. 10.1016/j.oraloncology.2012.06.019.

27. Cao W, Liu Z, Gokavarapu S, Chen Y, Yang R, Ji T. Reformed smokers have survival benefits after head and neck cancer. Br J Oral Maxillofac Surg. 2016;54(7):818–825. 10.1016/j.bjoms.2016.06.013.

28. Mayadev J, Lim J, Durbin-Johnson B, Valicenti R, Alvarez E. Smoking Decreases Survival in Locally Advanced Cervical Cancer Treated With Radiation. Am J Clin Oncol. 2018;41(3):295–301. 10.1097/COC.0000000000000268.

29. Dotto GP, Rustgi Anil K. Squamous Cell Cancers: A Unified Perspective on Biology and Genetics. Cancer Cell. 2016;29(5):622–637. 10.1016/j.ccell.2016.04.004.

30. Mulder FJ, Pierssens D, Baijens LWJ, Kremer B, Speel EM. Evidence for different molecular parameters in head and neck squamous cell carcinoma of nonsmokers and nondrinkers: Systematic review and meta-analysis on HPV, p16, and TP53. Head Neck. 2021;43(1):303–322. 10.1002/hed.26513.

31. Chen Y, Zhang Z, Jiang G, Zhao K. Gross tumor volume is the prognostic factor for squamous cell esophageal cancer patients treated with definitive radiotherapy. J Thorac Dis. 2016;8(6):1155–1161. 10.21037/jtd.2016.04.08.

32. Soliman M, Yaromina A, Appold S, et al. GTV differentially impacts locoregional control of non-small cell lung cancer (NSCLC) after different fractionation schedules: subgroup analysis of the prospective randomized CHARTWEL trial. Radiother Oncol. 2013;106(3):299–304. 10.1016/j.radonc.2012.12.008.

33. Linge A, Schmidt S, Lohaus F, et al. Independent validation of tumour volume, cancer stem cell markers and hypoxia-associated gene expressions for HNSCC after primary radiochemotherapy. Clin Transl Radiat Oncol. 2019;16:40–47. 10.1016/j.ctro.2019.03.002.

34. Sun C, Wang S, Ye W, et al. The Prognostic Value of Tumor Size, Volume and Tumor Volume Reduction Rate During Concurrent Chemoradiotherapy in Patients With Cervical Cancer. Front Oncol. 2022;12:934110. 10.3389/fonc.2022.934110.

35. Kather JN, Schulte J, Grabsch HI, et al. Deep learning detects virus presence in cancer histology. bioRxiv 2019. 10.1101/690206.

36. Klein S, Wuerdemann N, Demers I, et al. Predicting HPV association using deep learning and regular H&E stains allows granular stratification of oropharyngeal cancer patients. NPJ Digit Med. 2023;6(1):152. 10.1038/s41746-023-00901-z.

37. Wang R, Khurram SA, Walsh H, Young LS, Rajpoot N. A Novel Deep Learning Algorithm for Human Papillomavirus Infection Prediction in Head and Neck Cancers Using Routine Histology Images. Mod Pathol. 2023;36(12):100320. 10.1016/j.modpat.2023.100320.

38. Wang R, Gunesli GN, Skingen VE, et al. Deep learning for predicting prognostic consensus molecular subtypes in cervical cancer from histology images. NPJ Precis Oncol. 2025;9(1):11. 10.1038/s41698-024-00778-5.

39. MahmoodLab. UNI2-h; 2024. https://huggingface.co/MahmoodLab/UNI2-h.

40. Zimmermann E, Vorontsov E, Viret J, et al. Virchow2: Scaling Self-Supervised Mixed Magnification Models in Pathology. arxiv. 2024. 10.48550/arXiv.2408.00738.

41. Xu H, Usuyama N, Bagga J, et al. A whole-slide foundation model for digital pathology from real-world data. Nature. 2024;630(8015):181–188. 10.1038/s41586-024-07441-w.

42. Neidlinger P, Nahhas OSME, Muti HS, et al. Benchmarking foundation models as feature extractors for weakly-supervised computational pathology. arXiv. 2024. 10.48550/ARXIV.2408.15823.

43. Alfasly S, Alabtah G, Hemati S, Kalari KR, Garcia JJ, Tizhoosh HR. Validation of histopathology foundation models through whole slide image retrieval. Scientific Reports. 2025;15(1). 10.1038/s41598-025-88545-9.

44. Skripcak T, Belka C, Bosch W, et al. Creating a data exchange strategy for radiotherapy research: towards federated databases and anonymised public datasets. Radiother Oncol. 2014;113(3):303–309. 10.1016/j.radonc.2014.10.001.

